# WEDGE: imputation of gene expression values from single-cell RNA-seq datasets using biased matrix decomposition

**DOI:** 10.1101/864488

**Authors:** Yinlei Hu, Bin Li, Wen Zhang, Nianping Liu, Pengfei Cai, Falai Chen, Kun Qu

## Abstract

The low capture rate of expressed RNAs from single-cell sequencing technology is one of the major obstacles to downstream functional genomics analyses. Recently, a number of imputation methods have emerged for single-cell transcriptome data, however, recovering missing values in very sparse expression matrices remains a substantial challenge. Here, we propose a new algorithm, WEDGE (WEighted Decomposition of Gene Expression), to impute gene expression matrices by using a biased low-rank matrix decomposition method (bLRMD). WEDGE successfully recovered expression matrices, reproduced the cell-wise and gene-wise correlations, and improved the clustering of cells, performing impressively for applications with multiple cell type datasets with high dropout rates. Overall, this study demonstrates a potent approach for imputing sparse expression matrix data, and our WEDGE algorithm should help many researchers to more profitably explore the biological meanings embedded in their scRNA-seq datasets.

## INTRODUCTION

Single-cell sequencing technology has been widely used in studies of many biological systems, including but not limited to embryonic development (Xue et al. 2013; Yan et al. 2013; Klein et al. 2015), neuronal diversity (Zeisel et al. 2015; Lake et al. 2016; Lake et al. 2018) and a large variety of diseases (Zheng et al. 2017; Guo et al. 2018; Zhang et al. 2018; Peng et al. 2019a; Zhang et al. 2019). Despite the rapid increase in sequencing throughput, the number of captured genes per cell is still limited by technical challenges (Grün et al. 2014; Stegle et al. 2015; Kiselev et al. 2019; Hou et al. 2020). The dropout events in single-cell sequencing experiments lead to sparse expression matrices, which hinders subsequent studies (Kiselev et al. 2019; Tian et al. 2019). To overcome this, a variety of algorithms have been developed to impute the zero elements in the expression matrices (Chen and Zhou 2018; Huang et al. 2018; Linderman et al. 2018; van Dijk et al. 2018; Eraslan et al. 2019; Peng et al. 2019b; Wagner et al. 2019; Wang et al. 2019; Elyanow et al. 2020).

For example, MAGIC (van Dijk et al. 2018) recovers gene expression by using data diffusion to construct an affinity matrix which attempts to represent the neighborhood of similar cells. Huang *et al*. combined Bayesian and Poisson LASSO regression methods into SAVER (Huang et al. 2018) to estimate prior parameters and to restore missing elements of an expression data matrix, based on the assumption that gene expression follows the negative binomial distribution. Recently, they upgraded this approach to SAVER-X (Wang et al. 2019) by training a deep autoencoder model with gene expression patterns obtained from public single-cell data repositories. Eraslan *et al*. developed a deep neuron network model, DCA (Eraslan et al. 2019), which can denoise scRNA-seq data by learning gene-specific parameters. Many other tools have also emerged recently, such as SCRABBLE (Peng et al. 2019b), VIPER (Chen and Zhou 2018), ENHANCE (Wagner et al. 2019), ALRA (Linderman et al. 2018), and netNMF-sc (Elyanow et al. 2020), each of which seeks to improve recovery of the expression matrix for single-cell data. However, for datasets with high dropout rates— which therefore have very sparse expression matrices—it is still a challenge to abundantly recover gene expression data while avoiding over-imputing (Tian et al. 2019; Elyanow et al. 2020).

Here, we introduce a new algorithm, WEDGE, to impute gene expression values for sparse single-cell data based on low-rank matrix decomposition (Kim and Park 2008; Kim and Choi 2009; Wang et al. 2015). We applied WEDGE to multiple scRNA-seq datasets and compared its performance against several state-of-the-art methods. We also examined WEDGE’s ability to distinguish cell subpopulations in the Tabula Muris dataset (Tabula Muris et al. 2018), and assessed its performance for accurately imputing marker gene expression in a recently released dataset for peripheral blood mononuclear cells (PBMCs) from COVID-19 patients (Guo et al. 2020). Finally, we assessed the computer resources WEDGE consumes when analyzing large datasets.

## RESULTS

### Algorithm, performance, and robustness of WEDGE

The expression matrices obtained from single-cell sequencing experiments are sparse, caused by the low RNA capture rates during experimental sampling and processing (Chen and Zhou 2018; Eraslan et al. 2019). Weighted non-negative matrix factorization (WNMF) has demonstrated its potential for recovering missing elements from a sparse matrix (Kim and Choi 2009; Elyanow et al. 2020). However, it should be noted that the contribution of the zero elements in the raw matrix is completely ignored in the WNMF optimization process. In WEDGE, we adopted a low weight (0 ≤ λ ≤ 1) for the zero elements in the raw expression matrix during the low-rank decomposition (a so-called bLRMD method), and generated a convergent imputed matrix using an alternating non-negative least-squares algorithm (Fig. 1A and Methods). We chose not to set λ as zero, as the contribution of the zero elements is not completely negligible (Eraslan et al. 2019). Notably, as WEDGE is a completely unsupervised algorithm, it allows us to impute expression data matrices without any prior information about genes or cell types.

**Figure 1.**
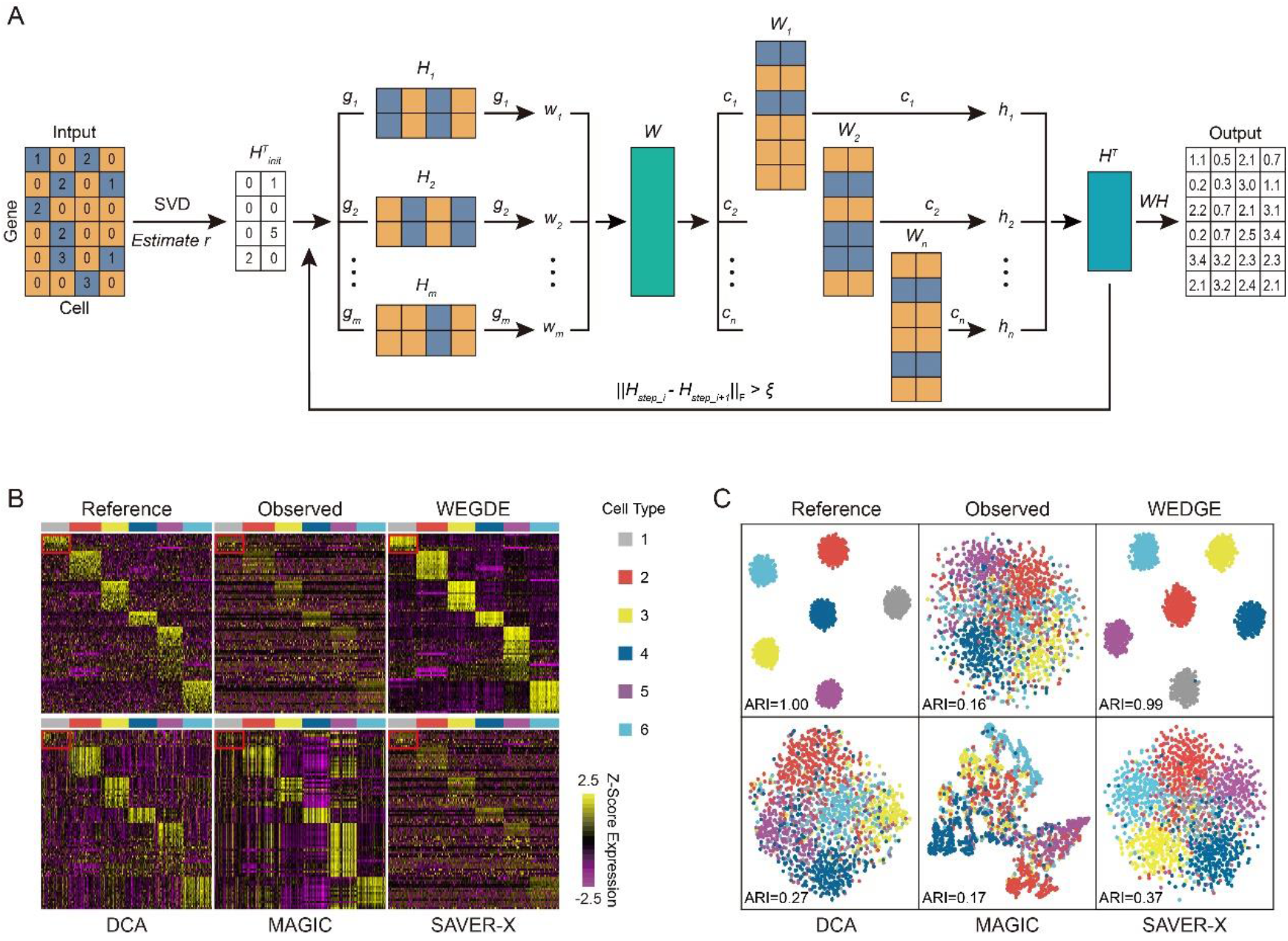
Design of the WEDGE algorithm and its performance for the simulated dataset. (**A**) Conceptual overview of the WEDGE workflow, where ***H*** and ***W*** are the lowdimensional representations of cells and genes respectively, ***g***_*i*_ is the *i*th gene, ***c**_j_* is the *j*th cell, ***w**_j_* is the *i*th row of ***W***, and ***h**_j_* is the *j*th column of ***H***. The yellow elements in ***H**_i_* and ***W**_j_* correspond to the zero elements in column *i* and row *j* of the original expression matrix, respectively, while the steel-blue elements represent the non-zero elements. is the convergence criterion (=1×10^-5^ by default). (**B**) Expression matrices of the top DE genes of the simulated reference and observed data (with dropout rate=0.45), and the results generated by WEDGE, DCA, MAGIC, and SAVER-X. The color bar at the top indicates different cell types. (**C**) tSNE (t-distributed Stochastic Neighbor Embedding) maps of the cells from the expression matrices imputed by different methods.

To test the performance of WEDGE in restoring gene expression, we first applied it to assess a simulated dataset generated using Splatter (Zappia et al. 2017) and compared WEDGE against DCA (Eraslan et al. 2019), MAGIC (van Dijk et al. 2018), and SAVER-X (Wang et al. 2019) (Fig. 1B). The reference data includes distinct marker genes for 6 different cell types and a dense expression matrix (original sparsity=10%, where sparsity is the percentage of zero elements). Based on the assumption that gene expression values follow a negative binomial distribution (Zappia et al. 2017), the Splatter simulation set 45% of the elements in the reference data to zero (*i.e*., dropout rate=0.45) to simulate dropout events and to obtain a down-sampled matrix, which we refer to as the “observed” data. The dropout events obscured the significance of the differentially expressed (DE) genes, but WEDGE successfully recovered their expression patterns, obtaining an imputed matrix apparently similar to the reference matrix, especially for the DE genes in cell type 1.

Also, we adopted the tSNE algorithm to explore the intercellular relationships in two-dimensional space, and used the adjusted random index (ARI) (Kiselev et al. 2017; Huang et al. 2018) to assess the accuracy of the cell clustering results (Fig. 1C), wherein a higher ARI value indicates that the clustering result is relatively closer to the “true” cell types. Using the expression matrix imputed by WEDGE, we can clearly distinguish these cell types. The ARI value of the cell clusters from the WEDGE imputed matrix is 0.99, higher than those from the other three imputation methods.

We further evaluated the robustness of WEDGE by applying it to impute “observed” matrices with different dropout rates (Supplemental Fig. S1). Interestingly, for the observed matrix with a low dropout rate (0.15), all the four methods (WEDGE, DCA, MAGIC, and SAVER-X) successfully recovered the distinctions between the cell types. However, for the data with higher sparsity (*i.e*., dropout rate>0.45) only WEDGE can still delineate the cell identities, demonstrating the advantage of WEDGE on imputing scRNA profiles with low capture rate.

In addition, to check whether the algorithm leads to over-imputing—for example, erroneously restoring non-DE genes so that they appear as DE genes—we applied WEDGE on another Splatter simulated dataset comprising 2 cell types (each with 1000 cells), 38 DE genes, and 162 non-DE genes. The sparsity of the reference and “observed” datasets were 7% and 40%, respectively. We clustered the cells using the DE gene expression values imputed by WEDGE, and the ARI value obtained was equal to 0.99, which was higher than other methods. (Supplemental Fig. S2A). In contrast, it was not possible to distinguish cell types based on the non-DE genes after WEDGE imputation (Supplemental Fig. S2B), implying that WEDGE is robust to over-imputing.

### Recovery performance for real scRNA-seq datasets

To examine the performance of WEDGE on real scRNA-seq data, we applied it to Zeisel’s dataset (Zeisel et al. 2015) on mouse brain scRNA-seq. We first constructed the reference matrix by extracting all the cells with more than 10000 UMI counts and all the genes detected in more than 40% of cells, and then generated an “observed” matrix with high sparsity by randomly setting a large proportion of the non-zero elements to zeros (dropout rate=0.85). From the heatmaps of gene expression matrices (Fig. 2A), we can see that WEDGE recovered the expression of the DE genes, especially those differentially expressed between interneurons and S1 pyramidal cells.

**Figure 2.**
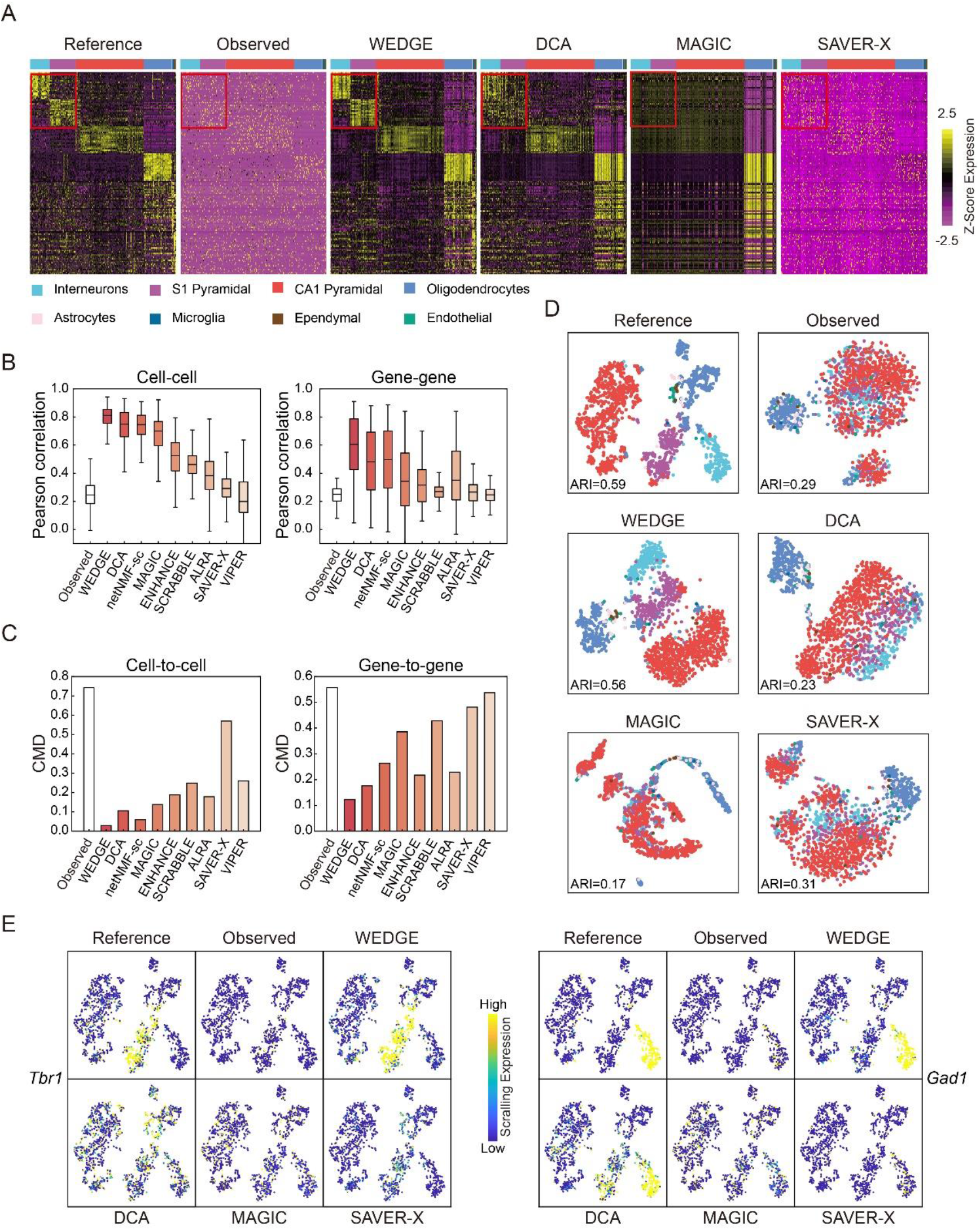
Application and performance assessment of WEDGE for Zeisel’s single-cell sequencing dataset, compared to existing methods. (**A**) Visualizations of the expression matrices of the top DE genes of different cell types, including the reference data, the observed data (dropout rate=0.85), and the imputed data generated by four different methods. The color bar at the top indicates known cell types. (**B**) Pearson correlation coefficients between the reference and imputed matrices for cells (left panel) and genes (right panel). Center line, median; box limits, upper and lower quartiles; whiskers, 1.5x interquartile range. (**C**) Distances of the cell-to-cell (left panel) and gene-to-gene (right panel) correlation matrices between the reference and imputed data. (**D**) 2-D tSNE maps of the cells from the reference, observed, and imputed datasets. The color scheme is the same as in (A). (**E**) Expression of the *Tba1* gene (left panel, a marker of S1 Pyramidal cells) and the *Gad1* gene (right panel, a marker of interneurons) as imputed by different methods, rendered in tSNE space.

We also used other tools, including SCRABBLE (Peng et al. 2019b), VIPER (Chen and Zhou 2018), ENHANCE (Wagner et al. 2019), ALRA (Linderman et al. 2018), and netNMF-sc (Elyanow et al. 2020) to assess the same “observed” matrix (Supplemental Fig. S3A). To quantify the similarity between the reference and imputed expression matrices, we calculated the cell-wise and gene-wise Pearson correlations between them (Huang et al. 2018), where higher correlation coefficients indicate better recovery performance. For cell-wise correlation coefficients, the WEDGE result (median value=0.81) is the highest among all the tested methods (Fig. 2B). The gene-wise correlation coefficients from WEDGE was also higher than that from the rest of the methods. Moreover, we computed the correlation matrix distances (CMDs) (Herdin et al. 2005; Huang et al. 2018) between the reference and imputed data, where a lower CMD indicates that the imputed data is closer to the reference data (Fig. 2C). For the matrix generated by WEDGE, the cell-to-cell CMD is 0.03 and the gene-to-gene CMD is 0.12, which are each tied for the lowest of all the tested methods. These comparisons together highlight that our WEDGE approach can recover both the cell-cell and genegene correlations from sparse single-cell RNA-seq datasets.

In the tSNE map of cells, WEDGE can clearly distinguish interneurons, S1 pyramidal neurons, and CA1 pyramidal neurons, and the ARI value of 0.56 for the clustering result calculated from its imputed matrix is the highest among all tested methods (Fig. 2D; Supplemental Fig. S3B). In particular, in visualizing the expression of an interneuron marker gene *Gad1* (Zeisel et al. 2015) and an S1 pyramidal marker gene *Tbr1*, WEDGE appropriately recovered their expression levels in the corresponding cell types, without overestimating their expression in other cell types (Fig. 2E; Supplemental Fig. S3C). Furthermore, we also applied WEDGE to Zeisel’s raw dataset (Zeisel et al. 2015), which includes 3005 cells and 19973 genes. After processing with WEDGE, the clustering performance index ARI increased from 0.42 (raw data) to 0.56, the highest among all tested methods (Supplemental Fig. S4).

As another example to confirm the utility of WEDGE, we applied the same procedures described above to Baron’s pancreas single-cell dataset (Baron et al. 2016), and compared WEDGE with other imputation methods. WEDGE recovered the expression of most of the DE genes, especially those of the ductal and activated stellate cells (Supplemental Fig. S5A). Similarly, the cell-wise and gene-wise Pearson correlation coefficients from WEDGE are both greater than those from any other tested methods, emphasizing its strong recovery performance (Supplemental Fig. S5B). Moreover, WEDGE’s cell-to-cell and gene-to-gene CMDs of, respectively, 0.02 and 0.11 were each the lowest for any of the tested methods (Supplemental Fig. S5C). Finally, in terms of cell clustering, WEDGE clearly classified alpha, beta, delta, ductal, acinar, and gamma cells, with an ARI value of 0.80, higher than those from all the other methods (Supplemental Fig. S5D).

### Classification of cell subpopulations

To test that if WEDGE can be applied for large datasets with multiple tissues and organs, we used it to process the recently released Tabula Muris dataset of mouse (10X Chromium sequencing data) (Tabula Muris et al. 2018). We used the k-nearest-neighbor graph based method in Seurat to cluster cells for both the raw and imputed data, and presented the results in tSNE space (Supplemental Fig. S6A&B). Among the 54 cell clusters generated from the raw data, 45 clusters have Jaccard index values greater than 0.5, indicating that they were also identified in the clustering of the WEDGE imputed data. (Supplemental Fig. S6C; see Methods). Notably, the WEDGE imputation improved the clustering resolution for some cell types: a main cluster of B cells (Supplemental Fig. S7) was classified into 4 sub-clusters (clusters 1, 3, 50, and 53; Fig. 3A), whereas these sub-clusters were not separated based on the raw data (Fig. 3B). Most cells in clusters 1, 3, 53 are from the spleen, while cluster 50 is dominated by B cells from lungs (Fig. 3C).

**Figure 3.**
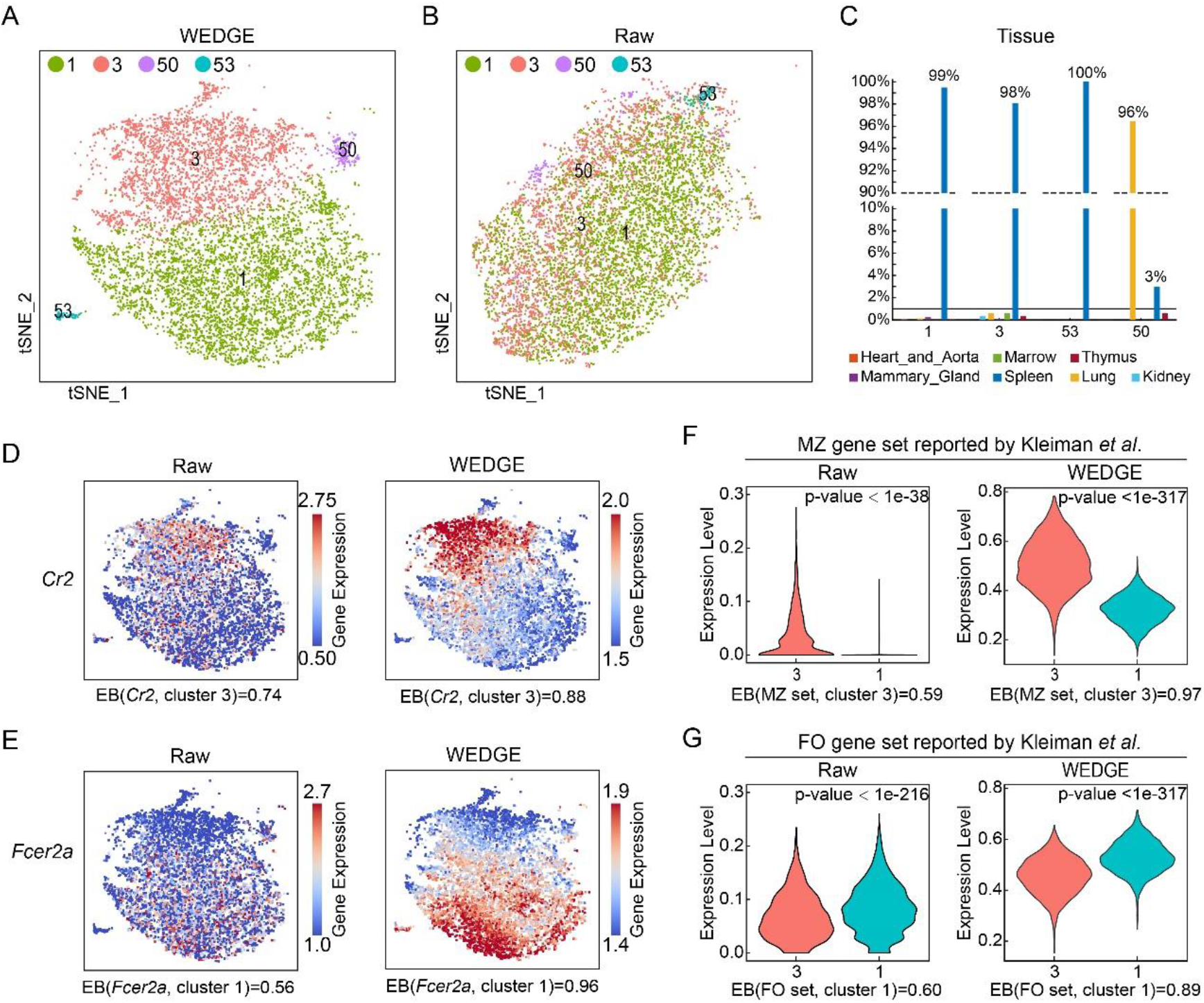
WEDGE imputation of the Tabula Muris dataset facilitated the classification of splenic B cell subpopulations. (**A, B**) 2-D tSNE maps of the splenic B cells generated from the WEDGE imputed data (A) and the raw data (B). The colors indicate cell clusters from the WEDGE imputed data. (**C**) The ratio of cells from different organs in splenic B cell clusters. (**D, E**) The expression of *Cr2* (D) or *Fcer2a* (E) in the raw and WEDGE imputed data, rendered in the 2-D tSNE space. EB: expression bias (see Methods). (**F, G**) The average expression of the marker gene sets of MZ (F) and FO (G) cells (reported by Kleiman *et al*. (Kleiman et al. 2015)) from the raw and WEDGE imputed data.

Cluster 3 splenic B cells strongly expressed the marginal zone B cell (MZ) marker *Cr2* (Martin and Kearney 2002), whereas cluster 1 splenic B cells were characterized by strong expression of the follicular B cell (FO) marker *Fcer2a* (Martin and Kearney 2002) (Fig. 3D&E). We developed a factor called “expression bias” (see Methods for the formula) to assess how a given imputation method affects the differential enrichment trend for inter-cluster expression comparisons. WEDGE imputation increased the expression bias of *Cr2* in cluster 3 from 0.74 to 0.88, and increased the expression bias of *Fcer2a* in cluster 1 from 0.56 to 0.96. The marker gene sets characteristic of MZ and FO splenic B cell subsets as reported by Kleiman (Kleiman et al. 2015) were also differentially expressed in cluster 3 and 1 respectively, and WEDGE respectively amplified the expression bias of these marker gene sets from 0.59/0.60 to 0.97/0.89 (Fig. 3F&G). Moreover, we tested the expression of the marker gene sets of the two cell types reported by Newman *et al*. (Newman et al. 2017), which also supported our classification of these two respective cell subpopulations as MZ and FO splenic B cells (Supplemental Fig. S8A&B). We noted that neither the raw data nor the WEDGE imputed data showed any obvious expression of the transitional B cell marker *Cd93* (Martin and Kearney 2002) (Supplemental Fig. S8C), and it was also notable that cluster 53 apparently represented an aggregation of cells with detected transcripts for fewer than 600 genes (Supplemental Fig. S8D).

We also used other state-of-the-art methods to impute the same dataset and checked their clustering results on the splenic B cells (Supplemental Fig. S9). DCA, MAGIC, ALRA, and ENHANCE enhanced the expression of *Cr2* and *Fcer2a* in some cells, but these methods also amplified batch effects, and clustering based on the imputation data from these methods did not clearly distinguish splenic B cells into FO and MZ subpopulations. SAVER-X did not classify the FO and MZ subpopulations based on differential expression trends for *Cr2* or *Fcer2a*. In addition, VIPER and SCRABBLE were unable to obtain imputation results from this dataset within 100 hours on the computer with 72 CPU-cores (2.2GHz) and 1TB memory, and netNMF-sc did not complete because of memory errors.

### Imputation of gene expression for the COVID-19 dataset

In a previous study of COVID-19 patients, Guo *et al*. (Guo et al. 2020) reported that one monocyte subset was found to strongly impact cytokine storms in patients classified as severe-stage. Cells of this severe-stage-specific monocyte subset strongly express many cytokines and related transcription factors such as *IL6, IL10, CXCL2, CXCL3, CCL4, ATF3, TNF*, and *HIVEP2*. However, dropout events in single cell sequencing experiments obscured this trend. Here, we applied WEDGE to this recently released COVID-19 dataset containing 13239 PBMCs from patients and 54951 PBMCs from healthy donors (Guo et al. 2020). The clusters obtained from the WEDGE imputed data correspond to cell clusters reported in the original paper (Guo et al. 2020), with an ARI value of 0.70, the highest among all tested methods (Fig. 4A&B). Guo *et al*. divided the monocyte cells into clusters 2, 9, 13, and 16, with the cells in cluster 9 apparently representing the severe-stage-specific monocyte subset (Guo et al. 2020). Following WEDGE imputation, the severe-stage-specific cytokines and upstream transcription factors reported by Guo *et al*. had increased expression bias in cluster 9 cells (from 0.03~0.93 to 1.00; Fig. 4D). Notably, the DE genes of cluster 9 generated from the WEDGE imputed data cover 99% of the DE genes in the raw data (Fig. 4C). The WEDGE imputed data also increased the expression bias of the *APOE* and *CXCR3* genes in cluster 9 cells (Supplemental Fig. S10A). Similar increases in expression bias were detected in the WEDGE imputed data for severe-stage-specific DE genes reported by Wilk *et al*. (Wilk et al. 2020) (Supplemental Fig. S10B).

**Figure 4.**
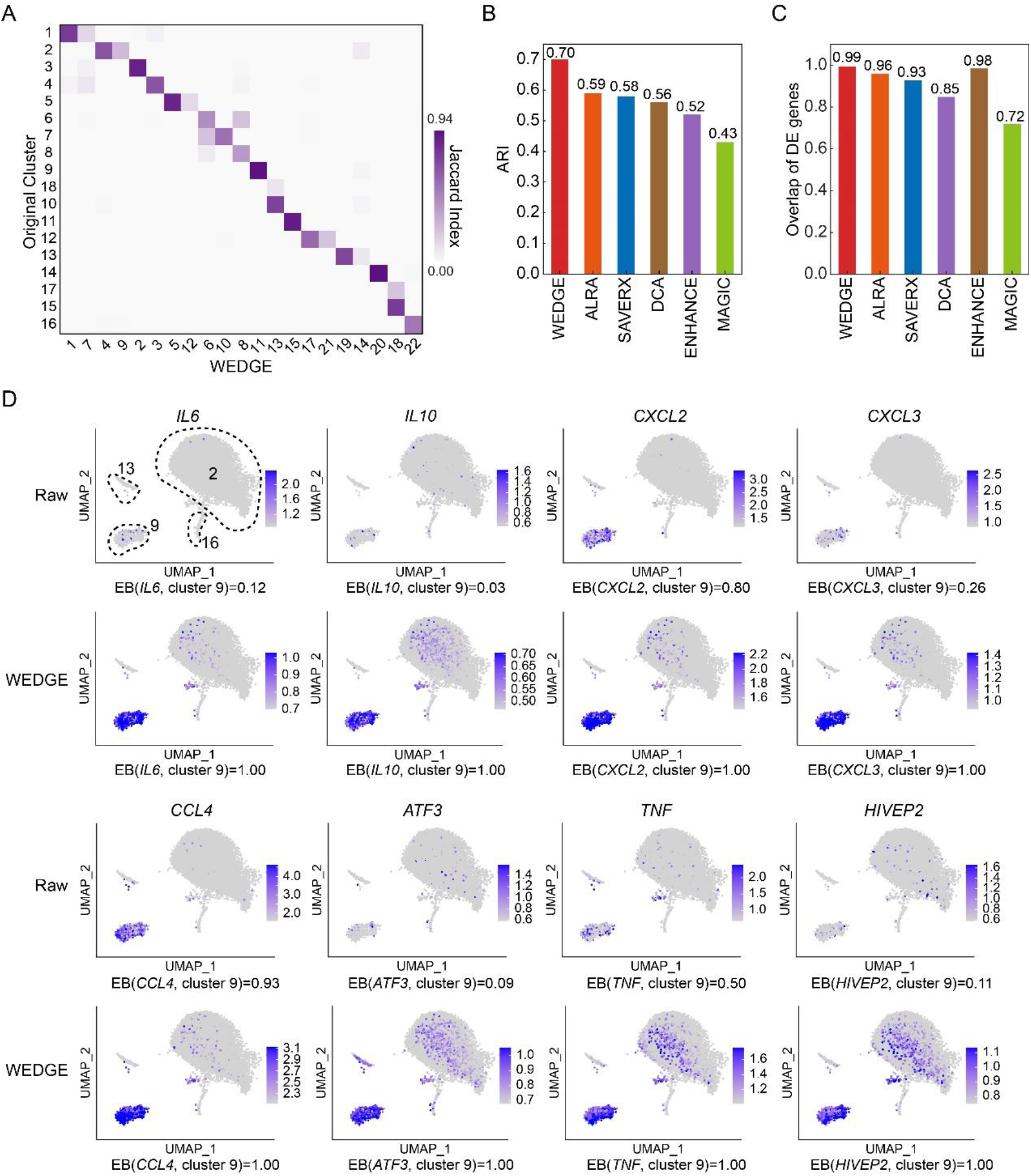
WEDGE enhanced marker gene expression for COVID-19 data from Guo *et al*. (**A**) Jaccard index between the cell clusters reported by Guo *et al*. (2020) and the clusters generated from WEDGE imputed data. (**B**) ARI values calculated from the clustering results of the imputed data generated by different methods. The results of other methods are not shown, as SCRABBLE did not finish the imputation within 100 hours on a computer with 72 CPU-cores (2.2GHz) and 1TB memory, and both VIPER and netNMF-sc reported memory errors. (**C**) Proportion of the reported DE genes that can be regenerated from the imputed data, *i.e*., 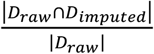, here, *D_raw_* and *D_imputed_* are DE gene sets generated from the raw data and the imputed data, respectively. (**D**) The expression of known marker genes of severe-stage specific monocytes. Clusters 2, 9, 13, and 16 are cell clusters reported by the original paper. EB: expression bias (see Methods).

### Scalability and efficiency

To quantify the scalability and efficiency of different imputation algorithms, we counted the time and memory consumption of WEDGE and other state-of-the-art methods when imputing Tabula Muris dataset that contains 55656 cells and 23341 genes (after filtering). WEDGE consumed 1 hour and up to 38GB of memory on a 72-core computer to complete the imputation process, which was close to the MAGIC method. (Fig. 5A&B). To further assess the computer resources that WEDGE spends on datasets of various sizes, we applied it to impute datasets comprising different numbers of cells (5000~1000000) but a fixed number of genes (2000), which were sampled from the mouse brain atlas project (see Methods). The runtime of WEDGE increased linearly with the number of cells, and its speed was close to DCA and MAGIC (Fig. 5C). For the dataset containing 1 million cells and 2000 genes, WEDGE finished the imputation of missing values in 12 minutes. Notably, WEDGE offers a visual interactive interface, making it convenient for researchers to use. We have uploaded WEDGE and the datasets used in this study to GitHub (https://github.com/QuKunLab/WEDGE).

**Figure 5.**
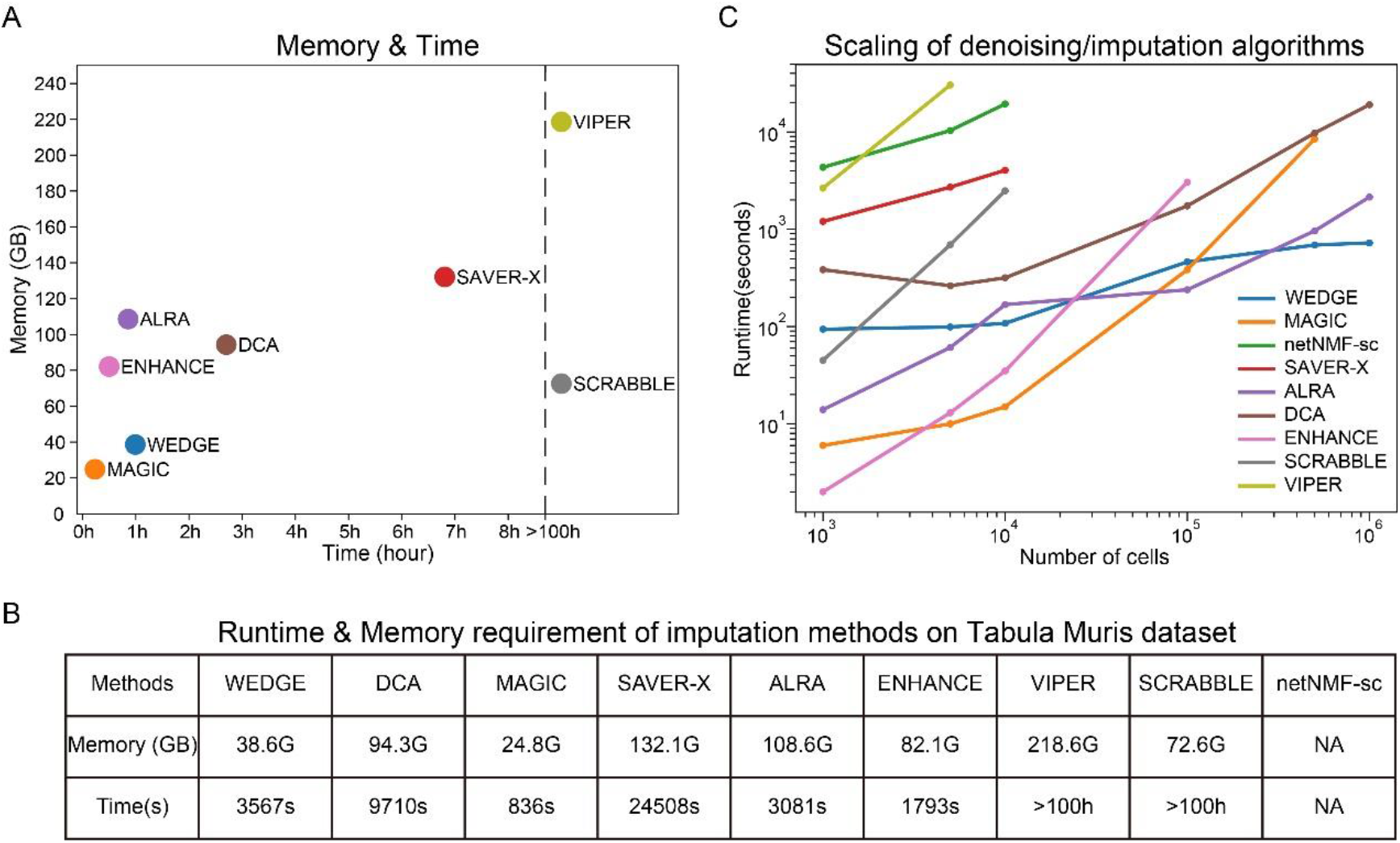
Computer resources consumed by different imputation methods, on a computer with 72 CPU-cores and 1TB memory. (**A, B**) The runtime and memory spent by different methods for imputing the Tabula Muris dataset, where netNMF-sc reported errors and stopped running. (**C**) The computational cost of different methods for imputing single-cell datasets of various sizes. Results that took more than 9 hours or reported memory errors are not shown here.

## DISCUSSION

Here, we present an approach, WEDGE, to impute missing gene expression information in single-cell sequencing datasets that is based on the combination of low-rank matrix decomposition and biased weight parameters for the zero and non-zero elements in the expression matrix. We demonstrate that the usage of WEDGE significantly improves the clustering accuracy of many scRNA-seq datasets, amplifies the contribution of differential genes to identifying cell types, and helps distinguish more cell subpopulations from low-quality data.

We tested the impact of the bias parameter *λ* on imputation, which indicated that recovery performance is insensitive to *λ* for datasets with dropout rates less than 0.6 (Supplemental Fig. S11; see Methods). For the datasets with dropout rates greater than 0.6, *λ* values between 0.1~0.15 can produce the best recovery results. We thus set *λ* = 0.15 for all datasets presented in this paper. The imputation contribution of the zero elements decreases with the increase of matrix sparsity, but it cannot be ignored, which implies that some zero elements may be related to the low expression of certain genes, rather than simply reflecting experimental noise.

There are still challenges for the informative imputation of scRNA-seq datasets, such as how to recover the heterogeneity between cell types instead of experimental batches, how to discover cell subtypes with very few cells from the imputed data, and how to use limited computer resources to process large datasets containing millions of cells. Moreover, it necessary to assess whether current imputation methods are applicable to datasets obtained using diverse bioanalytical methods beyond standard RNA-seq (e.g., single-cell ATAC-seq and profiling methods for various epigenomic modifications).

## METHODS

### Imputation model of WEDGE

WEDGE takes an expression matrix ***A***_*m*×*n*_ as input, where the element ***a***_*ij*_ represents the expression of the *i*th gene in the *j*th cell. By default, it normalizes the total expression of each cell to 10,000, and updates the expression value by performing a logarithm on it, *i.e*., 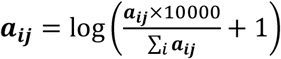. After imputation, WEDGE outputs two non-negative matrices, ***W*** and ***H***; we can then take their matrix product as the final imputed expression matrix, ***V***. In the WEDGE algorithm, we imputed single-cell sequencing data through the following optimization framework:

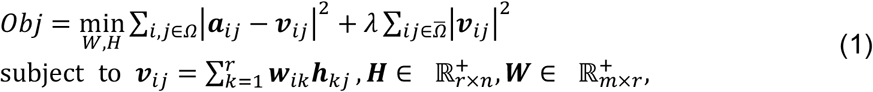

where 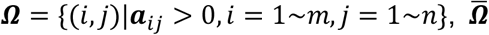 is the complementary set of ***Ω*** when the universe is {(*i,j*)|*i* = 1~*m,j* = 1~*n*}, ***v***_*ij*_ is the element of the *i*th row and *j*th column of ***V, w***_*ik*_ represents the element of matrix ***W***, and ***h***_*kj*_ is the element of matrix ***H***. In this model, the first term in the objective function guarantees minimization of approximation error between the non-zero elements of the original matrix ***A*** and the corresponding elements in the imputed matrix ***V***, while the second term tends to minimize the elements in ***V*** which correspond to zero elements in ***A***. The non-negative constraints in model (1) ensure that all the entries of ***V*** are no-negative.

Users can tune *λ* ∈ [0,1] to balance the contributions of the two terms of the objective functions. In order to study the influence of *λ* on the recovery performance, we down-sampled the reference data of the Zeisel and Baron datasets and adopted different dropout rates. By computing the correlation matrix distances (CMDs) (Herdin et al. 2005; Huang et al. 2018) between the reference data and the imputed data, we found that the recovery performance is not sensitive to the change of *λ* when the dropout rate is less than 0.6. For data with drop rate greater than 0.6, the best imputation result can be obtained by setting *λ* to 0.1~0.15. We set *λ* = 0.15 for all the datasets used in this study, which is also the default value for WEDGE.

### Optimization of the model

In the WEDGE algorithm, the matrix ***W*** and ***H*** were separately considered, which means that we fixed ***H*** to optimize ***W***, and then fixed ***W*** to generate the new ***H***. First, we defined that ***g***_*i*_ is the *i*th row of ***A***, 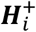 and 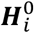 are composed of the ***H*** columns that correspond to the non-zero and zero elements of ***g***_*i*_ respectively, and 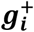 is the vector after deleting all zero elements from ***g***_i_. Then we rewrote the objective function of solving ***W***, as,

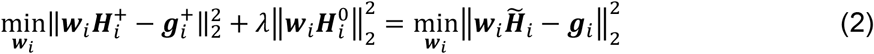

where 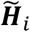 is the combination of 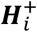 and 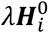 according to the original order of their elements in ***H***, and ***w***_*i*_ is the *i*th row of ***W***. In this case, optimizing ***W*** is equivalent to solving *m* non-negative least-squares problems (2) in parallel (Lawson and Hanson 1995). After ***W*** was obtained, we fixed it and solved ***H*** using similar algorithm as described above.

#### Algorithm 1. Optimization of WEDGE

~~~
Step1: generate the initial 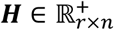 from singular value decomposition.
Step2: from a given ***H***, solve ***W*** in parallel with a non-negative least-square method.
Step3: from the ***W*** obtained in step 2, calculate a new ***H***.
Step4: iteratively return back to step 2 and 3 until the relative difference in the object function between two adjacent loops is less than 1×10^-5^ or the maximum specified number of iterations is reached.
~~~

### Estimating the rank of the expression matrix

During the optimization process for WEDGE, we designed a heuristic algorithm to determine the rank of ***A***_*m*×*n*_ based on the relative variation of its singular values (*σ_i_*: *i* = 1~min(*m,n*)}. For a descending list of *λ_i_*, we defined a function *f*(*σ_i_*) as

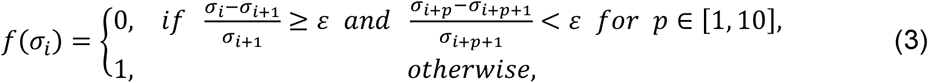

where *ε* is a small non-negative constant (0.085 by default). The following provides the details of the algorithm for the evaluation of rank *r*:

#### Algorithm 2: Estimating the rank of the expression matrix

~~~
Input: the singular values of matrix ***A*** : *σ*_1_ ≥ *σ*_2_ ≥ *σ*_3_ ≥ ⋯ ≥ *σ*_min(*m,n*)_
Output: the final rank retained for ***A*** : *r*
Algorithm:
   *r* = 1;
   while *f*(*σ_r_*) and *r* ≤ min(*m,n*) – 11 do
      *r* = *r* + 1;
   end while
~~~

### Correlation matrix distance (CMD)

CMD is usually used to determine the difference between two correlation matrices. It denotes the error of imputation process (Huang et al. 2018; Wang et al. 2019), as a larger CMD value indicates greater difference between the reference/raw matrix and the imputed matrix. The CMD of two correlation matrices ***R***_1_, ***R***_2_ is expressed as 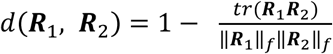.

### Jaccard index

The Jaccard index can be used to evaluate the similarity between two clusters (Levandowsky and Winter 1971). The Jaccard index of two clusters ***A, B*** is defined as 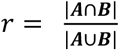, here |·| represented the number of elements in a set.

### Expression bias

The expression bias of gene *α* in cell cluster *C_i_* refers to the proportion of cells (in cluster *C_i_*) whose expression level of gene *α* is higher than the average expression of other clusters:

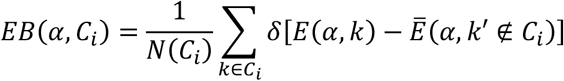

where *k* is a cell belonging to cluster *C_i_*, *E*(*α, k*) is the expression value of gene α in cell *k*, *k*′ is a cell belonging to other clusters, 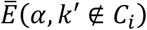 is the average expression value of gene α in all other clusters, *N*(*C_i_*) is the number of cells in cluster *C_i_*, and *δ* function is defined as

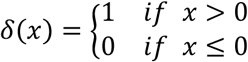

### Generation of the simulated scRNA-seq datasets

We used the *splatSimulate()* function in the Splatter R package (Zappia et al. 2017) to generate simulated datasets. For the dataset containing 6 cell types, 500 genes, and 2000 cells (shown in Fig. 1B-C and Supplemental Fig. S1), we set seed=42 and dropout.shape=-1, and its dropout rate was tuned by parameter dropout.mid values ranging from 1 to 6. For the dataset with two cell types, 200 genes, and 2000 cells (shown in Supplemental Fig. S2), we set seed=42, dropout.shape=-1, and dropout.mid=2.

### Processing of different datasets

For Zeisel’s dataset of the mouse cortex and hippocampus cells (GSE60361) (Zeisel et al. 2015), we generated a reference dataset that contains high quality cells and genes (performed identically with the previously described filtering step of SAVER (Huang et al. 2018)), and retained all of the marker genes described in the initial Zeisel study (*Tbr1, Spink8, Aldoc, Gad1, Mbp*, and *Thy1*). Then, we randomly set 85% of the non-zero elements of the reference data to zeros to generate observed data with dropouts. For Baron’s dataset of human pancreatic islet cells (GSE84133) (Baron et al. 2016), we also used the same process to filter the high quality cells and genes from the original data to build the reference dataset, in which 54% of the elements had non-zero values. We then randomly set 65% of the non-zero elements to zeros to simulate dropout events. For Tabula Muris dataset (GSE109774) (Tabula Muris et al. 2018), we applied the pipeline provided by Tabula Muris to filter the genes and cells, and obtained an expression matrix with 23341 genes and 55656 cells. For the COVID-19 dataset, we filtered genes and cells using the same method as Guo *et al* (Guo et al. 2020), to obtain an expression matrix containing 23,324 genes and 68,190 cells.

### Parameters for other imputation tools

(1) For the application to all datasets in the paper, DCA (version 0.2.2) was performed on the expression matrices with default parameters (type = ‘zinb-conddisp’, hiddensize = ‘64,32,64’ and learningrate = 0.001). (2) We ran SAVER-X (version1.0.0) on the expression matrices with default parameters. Specifically, we set “data.species = Others, use.pretrain = F” for the simulated datasets, “data.species = Mouse, use.pretrain = T, pretrained.weights.file = mouse_AdultBrain.hdf5, model.species = Mouse” for Zeisel’s dataset, and “data.species = Human, use.pretrain = T, pretrained.weights.file = human_Devbrain_nomanno.hdf5, model.species = Human” for Baron’s dataset. (3) MAGIC (Python version 1.5.5) was performed with normalized expression matrices (same as WEDGE) and default parameters (decay = 15, t = ‘auto’, and knn_dist = ‘euclidean’). Specifically, we set the numbers of principal components to 20 and the number of nearest neighbors to 15. (4) SCABBLE (MATLAB version) was performed on the library size normalized expression matrices with default parameters (parameter = [1,0,1e-4], nIter = 100). (5) VIPER (version 0.1.1) was performed on the expression matrices with default parameters (num = 0.8 times the number of genes, percentage.cutoff = 0.1, minbool = FALSE, and alpha = 0.5). (6) ALRA (RunALRA function in Seurat 3.0 (Butler et al. 2018; Stuart et al. 2019a)) was performed on the expression matrices with default parameters (k = NULL, q = 10). (7) ENHANCE was performed on the expression matrices with default parameters (max-components = 50). (8) netNMF-sc (version 2.0) was run on the expression matrices with default parameters and the outputs were used for downstream analysis. Specifically, we set dimensions = 20 and normalize = False. For the matrix preprocessing step of all methods, we adopted the algorithms recommended by their respective tutorials. For tools that did not describe the preprocessing algorithm, we used the same procedure as WEDGE.

### Parameters for cell clustering

We used Scanpy (Wolf et al. 2018) (version 1.4.0) and the default parameters (n_neighbors = 15 and n_pcs=50 in neighbors() function) to cluster cells for each dataset. Particularly, we set the “resolution” parameter (in louvain() function) to 1 for the first simulated dataset (shown in Fig. 1B-C and Supplemental Fig. S1), to 0.5 for the second simulated dataset (shown in Supplemental Fig. S2), and to 0.3 for both of the two real datasets (Zeisel’s and Baron’s datasets), to get the highest ARI value for the cell clustering of each reference matrix. For the clustering of the observed and imputed matrices of a given dataset, we adopted the same parameters as we used for the corresponding reference matrix. We used Seurat 2.3.0 (as the original paper) with parameters dims.use = 1:30 and resolution = 1.1, to cluster cells of the raw and the imputed expression matrices of Tabula Muris dataset (Tabula Muris et al. 2018). For the raw and imputed matrices of Guo’s dataset (Guo *et al*. 2020), we used Seurat 3.2.0 (as the original paper) with parameters dims.use = 1:50 and resolution = 0.5 to cluster cells.

### Scalability analysis

Scalability analysis was performed on a super computer with four Intel Xeon E7-8860 v4 2.20 GHz CPUs (72 cores in total) and 1TB memory. We down-sampled the mouse brain atlas dataset downloaded from 10X Genomics website (https://support.10xgenomics.com/single-cell-gene-expression/datasets/1.3.0/1M_neurons) to construct the benchmark datasets with different number of cells (from 1000 to 1000000). First, we filtered out the genes that were only expressed in three or fewer cells, and normalized the library size of the dataset. Then, we used the gene filtering function of Scanpy, *i.e*., scanpy.pp.highly_variable_genes(), with min_mean=0.0125, max_mean=3, min_disp=0.5 and n_top_genes=2000 to obtain the top 2000 most variable genes. With the fixed number of genes, we sampled 1000, 5000, 10000, 100000, 500000, and 1000000 cells from the raw dataset to simulate experiments of different scales.

### TSNE, UMAP, and heatmap visualization

(1) Settings for dimension reduction: For the simulated dataset, Baron’s dataset, and Zeisel’s datasets, we used the first 20 principal components to perform tSNE analysis. For Tabula Muris dataset (Tabula Muris et al. 2018), we performed the 2D tSNE embedding by using Seurat with parameters dims.use = 1:30, seed.use = 10, and perplexity = 30. For Guo’s COVID-19 dataset, we used the same 2D UMAP method as the original paper (Guo et al. 2020). (2) Settings for heatmap: we calculated the Z-score of the expression of each gene, and used Seurat 3.0 (Butler et al. 2018; Stuart et al. 2019a; Stuart et al. 2019b) to extract the top DE genes from the reference matrix. The top 20 DE genes (only.pos = TRUE, min.pct = 0.1, and logfc.threshold = 0.25 in FindAllMarkers() function) of each cell type are shown in the heatmaps for Baron’s and Zeisel’s datasets (*i.e*., Fig. 2A; Supplemental Fig. S3A & 5A), and the top 30 DE genes (only.pos = TRUE, min.pct = 0.1, and logfc.threshold = 0.15 in FindAllMarkers() function) for each cell type are shown in the heatmaps of a simulated dataset (*i.e*., Fig. 1B). For the observed and imputed matrices of each dataset, we present the same DE genes in the same order as in the heatmap of the reference matrix.

## DATA ACCESS

Published datasets used in this study: Zeisel’s dataset of the mouse cortex and hippocampus cells (GSE60361), Baron’s dataset of human pancreatic islet cells (GSE84133) (Baron et al. 2016), Tabula Muris dataset including 20 mouse organs (GSE109774) (Tabula Muris et al. 2018), Guo’s dataset (Guo et al. 2020) containing PBMCs from 2 COVID-19 patients (GSE150861), and the 10X dataset of 2 healthy donors (https://support.10xgenomics.com/single-cell-gene-expression/datasets/3.1.0/5k_pbmc_NGSC3_aggr). The source code of WEDGE was released at https://github.com/QuKunLab/WEDGE.

## ACKNOWLEDGMENTS

This work was supported by the National Key R&D Program of China (2017YFA0102900 to K.Q.), the National Natural Science Foundation of China grants (91940306, 81788101, 31970858, 31771428 and 91640113 to K.Q.; 61972368, 11571338 to F.C.), and the Fundamental Research Funds for the Central Universities (YD2070002019 and WK2070000158 to K.Q.). We thank the USTC supercomputing center and the School of Life Science Bioinformatics Center for providing computing resources for this project.

## AUTHOR CONTRIBUTIONS

K.Q. and F.C. conceived and supervised the project; Y.H. and B.L. designed implemented, and validated WEDGE with the help from W.Z., N.L. and P.C.; B.L., Y.H. and K.Q. wrote the manuscript with inputs from all the authors.

## REFERENCES

Baron M, Veres A, Wolock SL, Faust AL, Gaujoux R, Vetere A, Ryu JH, Wagner BK, Shen-Orr SS, Klein AM et al. 2016. A Single-Cell Transcriptomic Map of the Human and Mouse Pancreas Reveals Inter- and Intra-cell Population Structure. Cell Syst 3: 346–360 e344.

Butler A, Hoffman P, Smibert P, Papalexi E, Satija R. 2018. Integrating single-cell transcriptomic data across different conditions, technologies, and species. Nature Biotechnology 36: 411.

Chen M, Zhou X. 2018. VIPER: variability-preserving imputation for accurate gene expression recovery in single-cell RNA sequencing studies. Genome Biol 19: 196.

Elyanow R, Dumitrascu B, Engelhardt BE, Raphael BJ. 2020. netNMF-sc: leveraging gene-gene interactions for imputation and dimensionality reduction in single-cell expression analysis. Genome research 30: 195–204.

Eraslan G, Simon LM, Mircea M, Mueller NS, Theis FJ. 2019. Single-cell RNA-seq denoising using a deep count autoencoder. Nat Commun 10: 390.

Grün D, Kester L, Van Oudenaarden A. 2014. Validation of noise models for single-cell transcriptomics. Nature methods 11: 637.

Guo C, Li B, Ma H, Wang X, Cai P, Yu Q, Zhu L, Jin L, Jiang C, Fang J. 2020. Single-cell analysis of two severe COVID-19 patients reveals a monocyte-associated and tocilizumab-responding cytokine storm. Nature Communications 11: 1–11.

Guo X, Zhang Y, Zheng L, Zheng C, Song J, Zhang Q, Kang B, Liu Z, Jin L, Xing R. 2018. Global characterization of T cells in non-small-cell lung cancer by single-cell sequencing. Nature medicine 24: 978.

Herdin M, Czink N, Ozcelik H, Bonek E. 2005. Correlation matrix distance, a meaningful measure for evaluation of non-stationary MIMO channels. In 2005IEEE61st Vehicular Technology Conference, Vol 1, pp. 136–140. IEEE.

Hou W, Ji Z, Ji H, Hicks SC. 2020. A systematic evaluation of single-cell RNA-sequencing imputation methods. Genome Biology 21: 218.

Huang M, Wang J, Torre E, Dueck H, Shaffer S, Bonasio R, Murray JI, Raj A, Li M, Zhang NR. 2018. SAVER: gene expression recovery for single-cell RNA sequencing. Nat Methods 15: 539–542.

Kim H, Park H. 2008. Nonnegative matrix factorization based on alternating nonnegativity constrained least squares and active set method. SIAM journal on matrix analysis and applications 30: 713–730.

Kim Y-D, Choi S. 2009. Weighted nonnegative matrix factorization. In 2009 IEEE International Conference on Acoustics, Speech and Signal Processing, pp. 1541–1544. IEEE.

Kiselev VY, Andrews TS, Hemberg M. 2019. Challenges in unsupervised clustering of single-cell RNA-seq data. Nat Rev Genet 20: 273–282.

Kiselev VY, Kirschner K, Schaub MT, Andrews T, Yiu A, Chandra T, Natarajan KN, Reik W, Barahona M, Green AR et al. 2017. SC3: consensus clustering of single-cell RNA-seq data. Nature Methods 14: 483.

Kleiman E, Salyakina D, De Heusch M, Hoek KL, Llanes JM, Castro I, Wright JA, Clark ES, Dykxhoorn DM, Capobianco E et al. 2015. Distinct Transcriptomic Features are Associated with Transitional and Mature B-Cell Populations in the Mouse Spleen. Front Immunol 6: 30.

Klein AM, Mazutis L, Akartuna I, Tallapragada N, Veres A, Li V, Peshkin L, Weitz DA, Kirschner MW. 2015. Droplet barcoding for single-cell transcriptomics applied to embryonic stem cells. Cell 161: 1187–1201.

Lake BB, Ai R, Kaeser GE, Salathia NS, Yung YC, Liu R, Wildberg A, Gao D, Fung H-L, Chen S. 2016. Neuronal subtypes and diversity revealed by single-nucleus RNA sequencing of the human brain. Science 352: 1586–1590.

Lake BB, Chen S, Sos BC, Fan J, Kaeser GE, Yung YC, Duong TE, Gao D, Chun J, Kharchenko PV. 2018. Integrative single-cell analysis of transcriptional and epigenetic states in the human adult brain. Nature biotechnology 36: 70–80.

Lawson CL, Hanson RJ. 1995. Solving least squares problems. Siam.

Levandowsky M, Winter D. 1971. Distance between Sets. Nature 234: 34–35.

Linderman GC, Zhao J, Kluger Y. 2018. Zero-preserving imputation of scRNA-seq data using low-rank approximation. bioRxiv doi:https://doi.org/10.1101/397588.

Martin F, Kearney JF. 2002. Marginal-zone B cells. Nat Rev Immunol 2: 323–335.

Newman R, Ahlfors H, Saveliev A, Galloway A, Hodson DJ, Williams R, Besra GS, Cook CN, Cunningham AF, Bell SE et al. 2017. Maintenance of the marginal-zone B cell compartment specifically requires the RNA-binding protein ZFP36L1. Nat Immunol 18: 683–693.

Peng J, Sun B-F, Chen C-Y, Zhou J-Y, Chen Y-S, Chen H, Liu L, Huang D, Jiang J, Cui G-S. 2019a. Single-cell RNA-seq highlights intra-tumoral heterogeneity and malignant progression in pancreatic ductal adenocarcinoma. Cell research 29: 725–738.

Peng T, Zhu Q, Yin P, Tan K. 2019b. SCRABBLE: single-cell RNA-seq imputation constrained by bulk RNA-seq data. Genome Biol 20: 88.

Stegle O, Teichmann SA, Marioni JC. 2015. Computational and analytical challenges in single-cell transcriptomics. Nature Reviews Genetics 16: 133.

Stuart T, Butler A, Hoffman P, Hafemeister C, Papalexi E, Mauck III WM, Hao Y, Stoeckius M, Smibert P, Satija R. 2019a. Comprehensive integration of single-cell data. Cell 177: 1888–1902. e1821.

Stuart T, Butler A, Hoffman P, Hafemeister C, Papalexi E, Mauck III WM, Hao Y, Stoeckius M, Smibert P, Satija R. 2019b. Comprehensive Integration of Single-Cell Data. Cell.

Tabula Muris C, Overall c, Logistical c, Organ c, processing, Library p, sequencing, Computational data a, Cell type a, Writing g et al. 2018. Single-cell transcriptomics of 20 mouse organs creates a Tabula Muris. Nature 562: 367–372.

Tian L, Dong X, Freytag S, Le Cao KA, Su S, JalalAbadi A, Amann-Zalcenstein D, Weber TS, Seidi A, Jabbari JS et al. 2019. Benchmarking single cell RNA-sequencing analysis pipelines using mixture control experiments. Nat Methods 16: 479–487.

van Dijk D, Sharma R, Nainys J, Yim K, Kathail P, Carr AJ, Burdziak C, Moon KR, Chaffer CL, Pattabiraman D et al. 2018. Recovering Gene Interactions from Single-Cell Data Using Data Diffusion. Cell 174: 716–729 e727.

Wagner F, Barkley D, Yanai I. 2019. Accurate denoising of single-cell RNA-Seq data using unbiased principal component analysis. BioRxiv doi:https://doi.org/10.1101/655365.

Wang J, Agarwal D, Huang M, Hu G, Zhou Z, Ye C, Zhang NR. 2019. Data denoising with transfer learning in single-cell transcriptomics. Nature Methods 16: 875–878.

Wang Z, Lai M-J, Lu Z, Fan W, Davulcu H, Ye J. 2015. Orthogonal rank-one matrix pursuit for low rank matrix completion. SIAM Journal on Scientific Computing 37: A488–A514.

Wilk AJ, Rustagi A, Zhao NQ, Roque J, Martínez-Colón GJ, McKechnie JL, Ivison GT, Ranganath T, Vergara R, Hollis T. 2020. A single-cell atlas of the peripheral immune response in patients with severe COVID-19. Nature Medicine. 1–7.

Wolf FA, Angerer P, Theis FJ. 2018. SCANPY: large-scale single-cell gene expression data analysis. Genome Biol 19: 15.

Xue Z, Huang K, Cai C, Cai L, Jiang C-y, Feng Y, Liu Z, Zeng Q, Cheng L, Sun YE. 2013. Genetic programs in human and mouse early embryos revealed by single-cell RNA sequencing. Nature 500: 593–597.

Yan L, Yang M, Guo H, Yang L, Wu J, Li R, Liu P, Lian Y, Zheng X, Yan J et al. 2013. Single-cell RNA-Seq profiling of human preimplantation embryos and embryonic stem cells. Nat Struct Mol Biol 20: 1131–1139.

Zappia L, Phipson B, Oshlack A. 2017. Splatter: simulation of single-cell RNA sequencing data. Genome Biol 18: 174.

Zeisel A, Muñoz-Manchado AB, Codeluppi S, Lönnerberg P, La Manno G, Juréus A, Marques S, Munguba H, He L, Betsholtz C. 2015. Cell types in the mouse cortex and hippocampus revealed by single-cell RNA-seq. Science 347: 1138–1142.

Zhang L, Yu X, Zheng L, Zhang Y, Li Y, Fang Q, Gao R, Kang B, Zhang Q, Huang JY. 2018. Lineage tracking reveals dynamic relationships of T cells in colorectal cancer. Nature 564: 268.

Zhang P, Yang M, Zhang Y, Xiao S, Lai X, Tan A, Du S, Li S. 2019. Dissecting the Single-Cell Transcriptome Network Underlying Gastric Premalignant Lesions and Early Gastric Cancer. Cell reports 27: 1934–1947. e1935.

Zheng C, Zheng L, Yoo J-K, Guo H, Zhang Y, Guo X, Kang B, Hu R, Huang JY, Zhang Q. 2017. Landscape of infiltrating T cells in liver cancer revealed by single-cell sequencing. Cell 169: 1342–1356. e1316.

