## Supplemental Fig for "WEDGE: imputation of gene expression values from single-cell RNA-seq datasets using biased matrix decomposition"

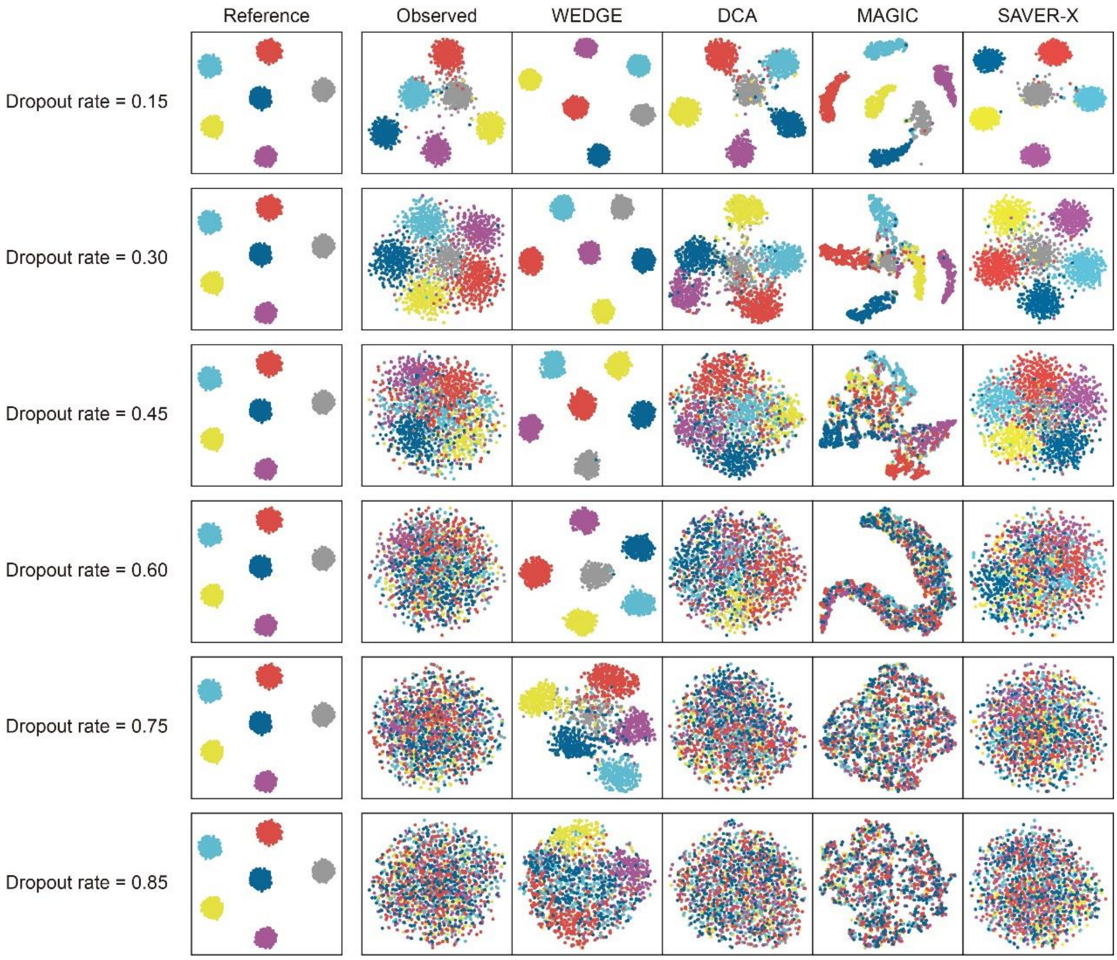


**Supplemental Figure S1.** Performance of different imputation algorithms for the recovery of the observed data with different dropout rates, visualized in tSNE space.

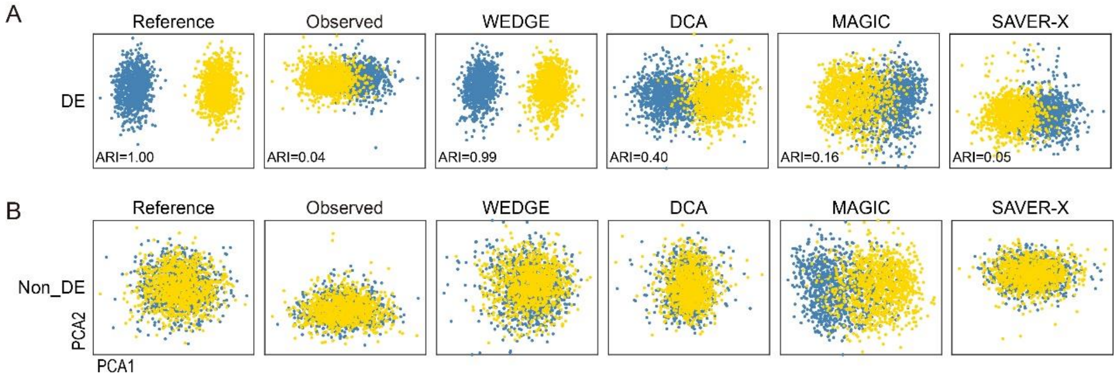


**Supplemental Figure S2.** Two-dimension PCA visualization of the cells from the reference data, observed data, and data imputed with 4 different methods. (**A**) Scores plots of PCA results for cells, calculated using expression data for the cell-type-specific DE genes (as defined from the reference dataset), based on the data imputed using four different imputation methods. (**B**) PCA results calculated using the imputed expression data for the non-DE genes.

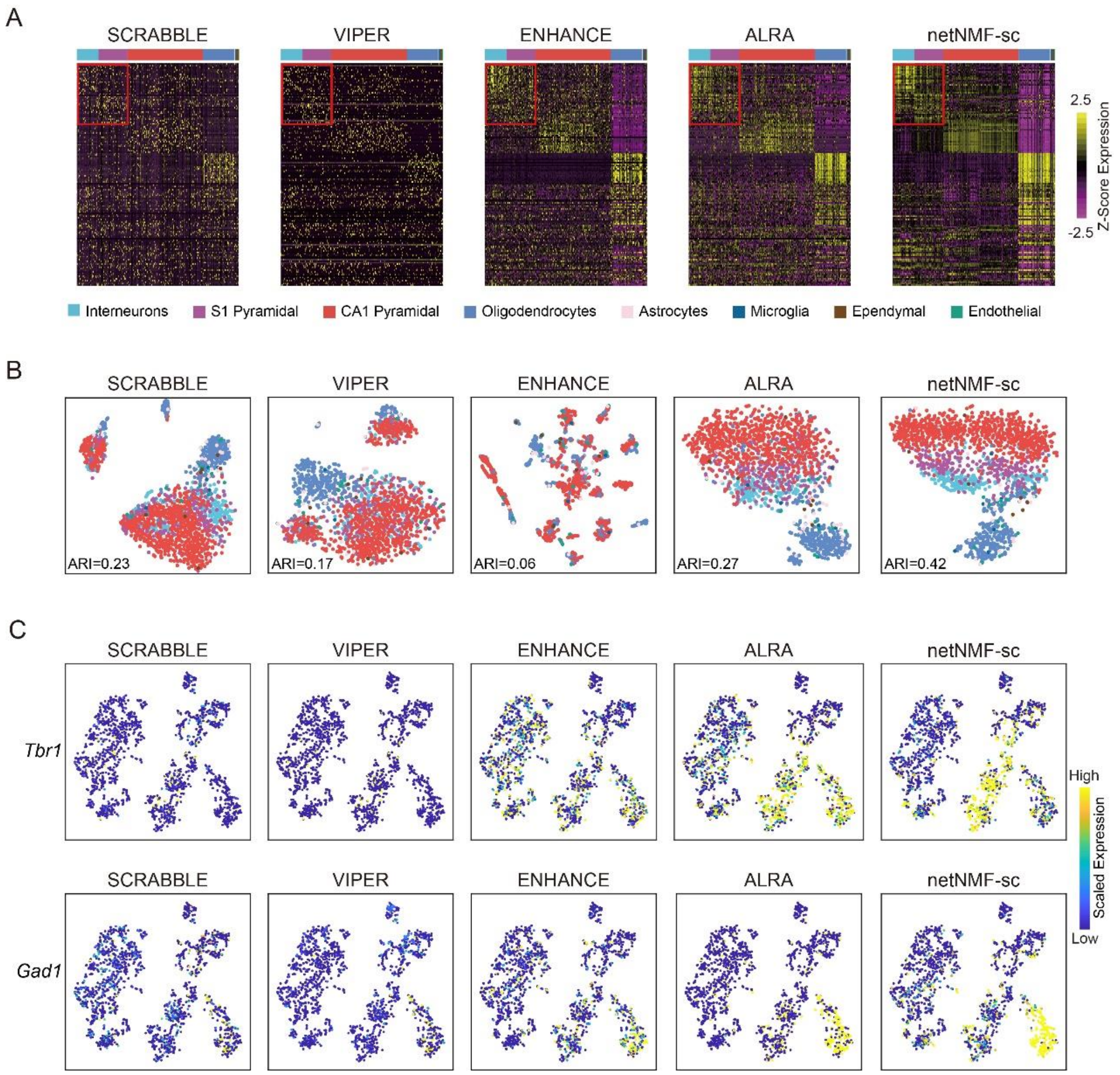


**Supplemental Figure S3.** The performance of SCRABBLE, VIPER, ENHANCE, ALRA, and netNMF-sc on Zeisel's dataset. (**A**) Visualization of the expression matrices of the top DE genes of different cell types, for the reference data, the observed data (dropout rate=0.85), and the imputed data generated by different methods. The color bar at the top indicates known cell types. (**B**) 2-D tSNE maps of the cells from the reference, observed, and variously imputed data. The color scheme is the same as in (A). (**C**) Expression of *Tba1* (a marker of S1 Pyramidal cells) and *Gad1* (a marker of interneurons), rendered in tSNE space.

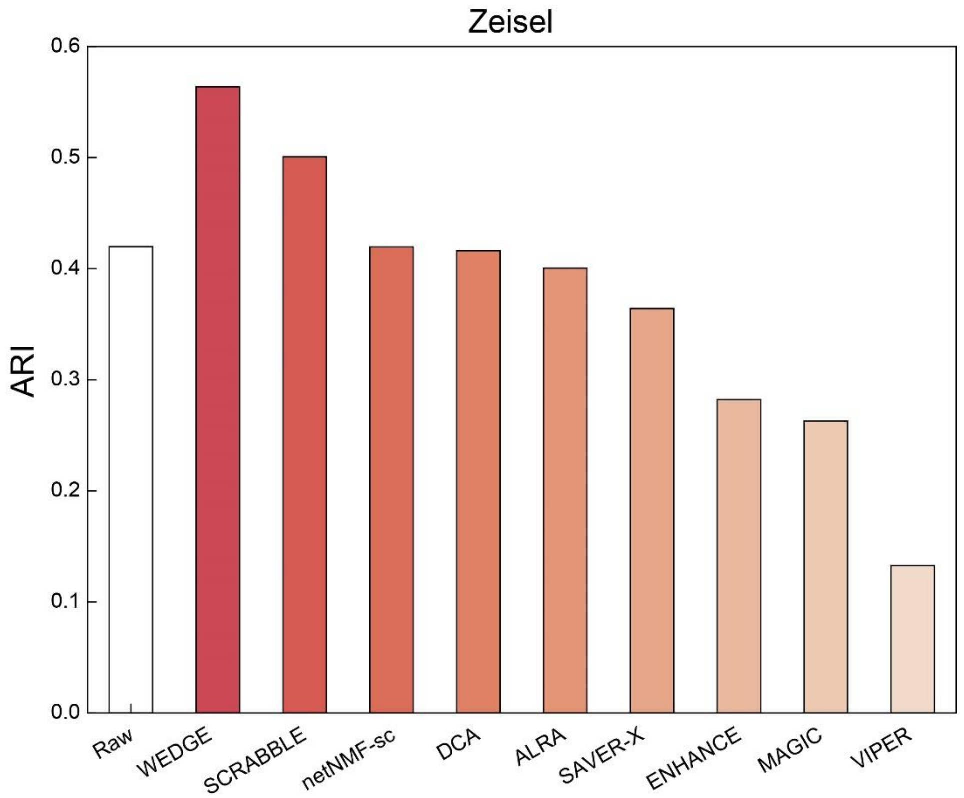


**Supplemental Figure S4.** The ARI values calculated from the clustering results of Zeisel's raw data and the imputed data generated by different methods.

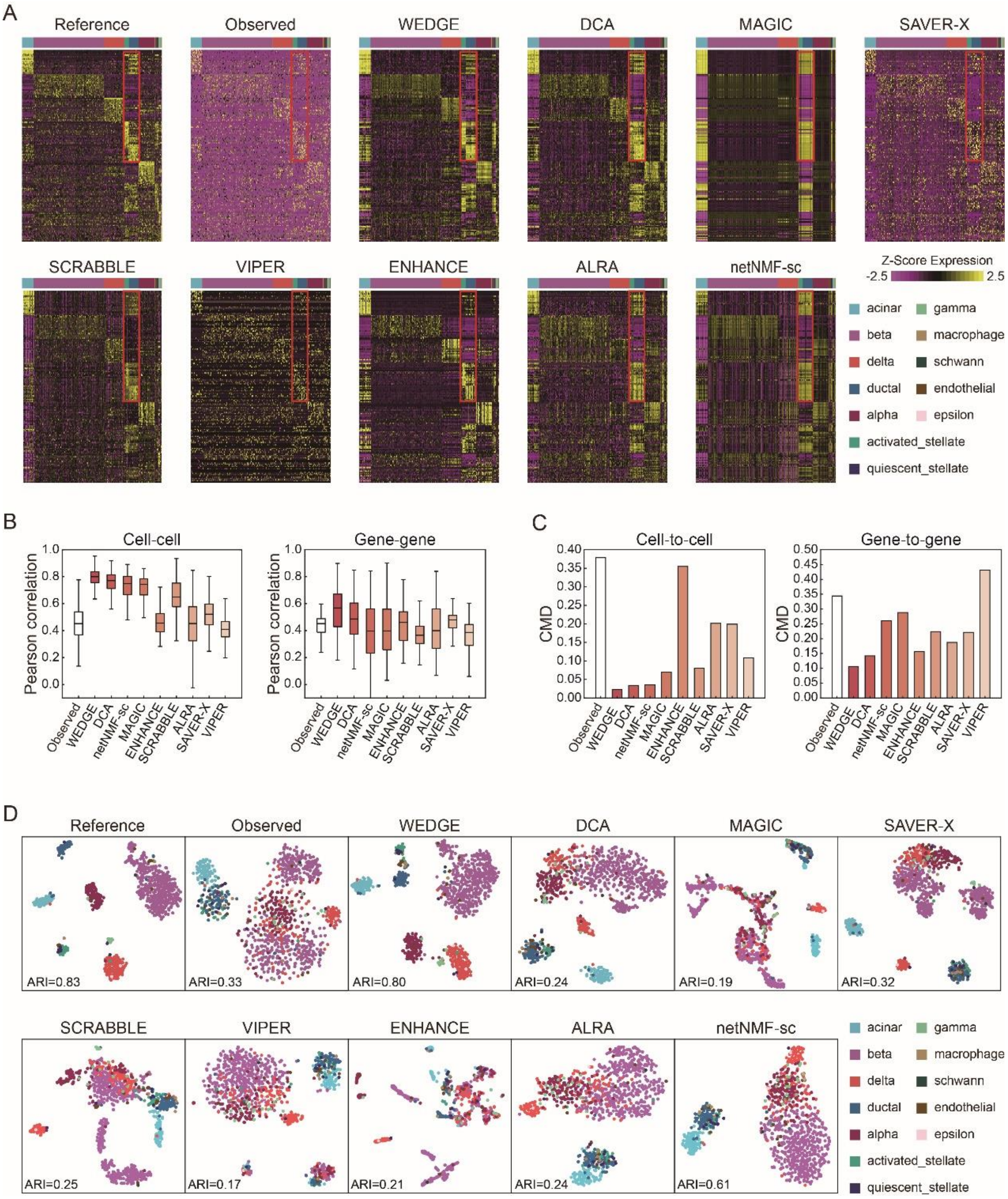

**Supplemental Figure S5.** Application and performance of WEDGE to Baron's single-cell sequencing dataset, compared to existing methods. (**A**) Visualization of the expression matrices of the top DE genes of different cell types, for the reference data, the observed data (dropout rate=0.65), and the imputed data generated by different methods. The color bar at the top indicates known cell types. (**B**) Pearson correlation coefficients between the reference and imputed matrices (as shown in (A)) for cells (left panel) and genes (right panel). Center line, median; box limits, upper and lower quartiles; whiskers, 1.5x interquartile range. (**C**) Distances of the cell-to-cell (left panel)

and gene-to-gene (right panel) correlation matrices between the reference and imputed datasets. (**D**) 2-D tSNE maps of the cells from the reference, observed, and variously imputed data. The color scheme is the same as in (A).

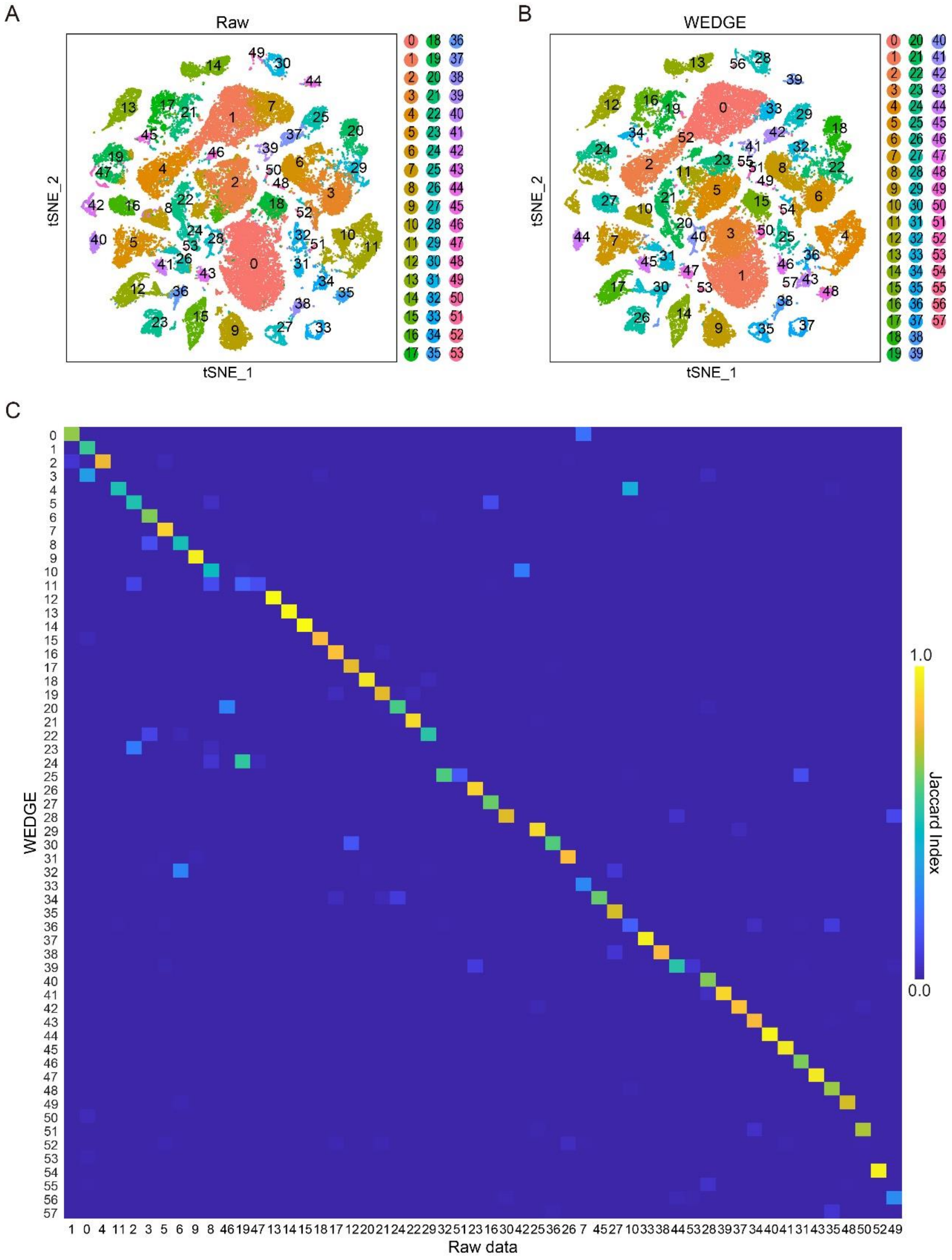

**Supplemental Figure S6.** Clustering of Tabula Muris Mouse Cell Atlas dataset. (**A**, **B**) 2-D tSNE maps of cells from the WEDGE imputed data. The colors denote the cell clusters generated from the raw data (A) and the WEDGE imputed data (B) respectively. (**C**) Jaccard index between cell clusters of the raw data and the WEDGE imputed data.

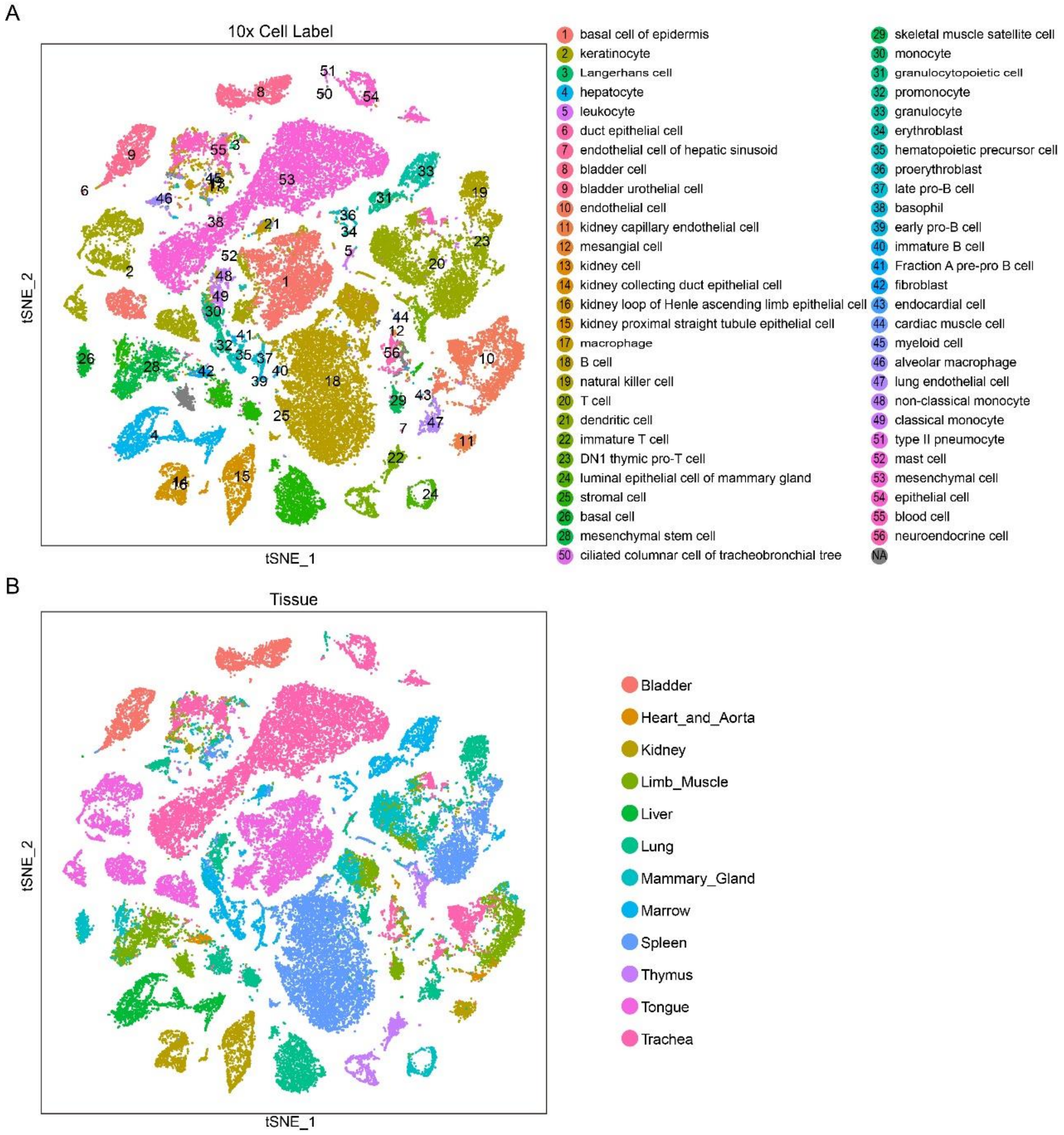


**Supplemental Figure S7.** Source information of cells in Tabula Muris Mouse Cell Atlas dataset. The colors indicate cell type (**A**) and organ type (**B**) reported by the Tabula Muris Consortium.

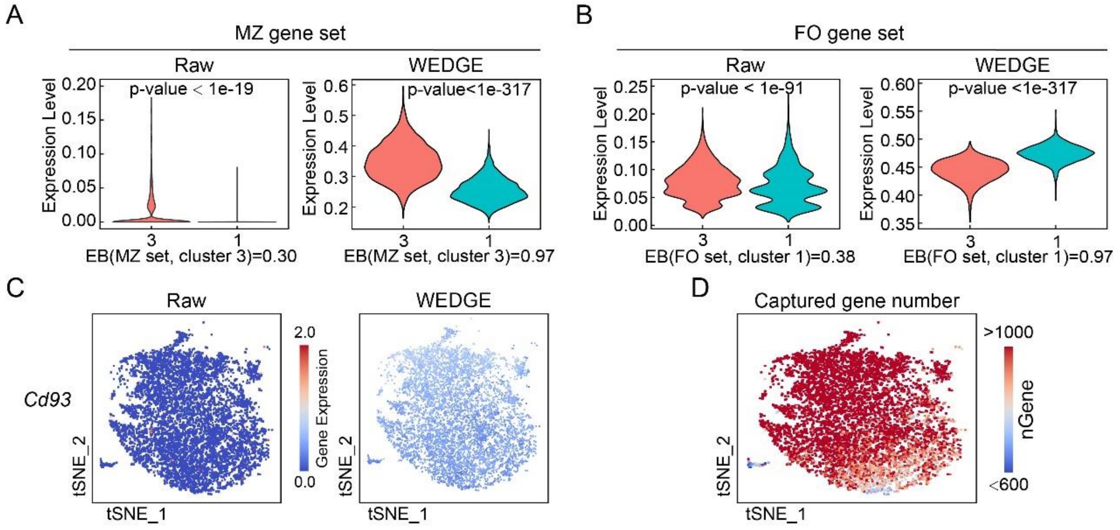


**Supplemental Figure S8.** Genomic features of splenic B cell subpopulations obtained from Tabula Muris dataset. (**A, B**) The average expression of the marker gene sets of MZ (A) and FO (B) cells (reported by Newman *et al*. [37]) from the raw (A) and WEDGE imputed (B) data. EB: expression bias (see Methods). (**C**) The raw and WEDGE imputed expression values of *Cd93* in splenic B cells. (**D**) The number of genes detected in each cell.

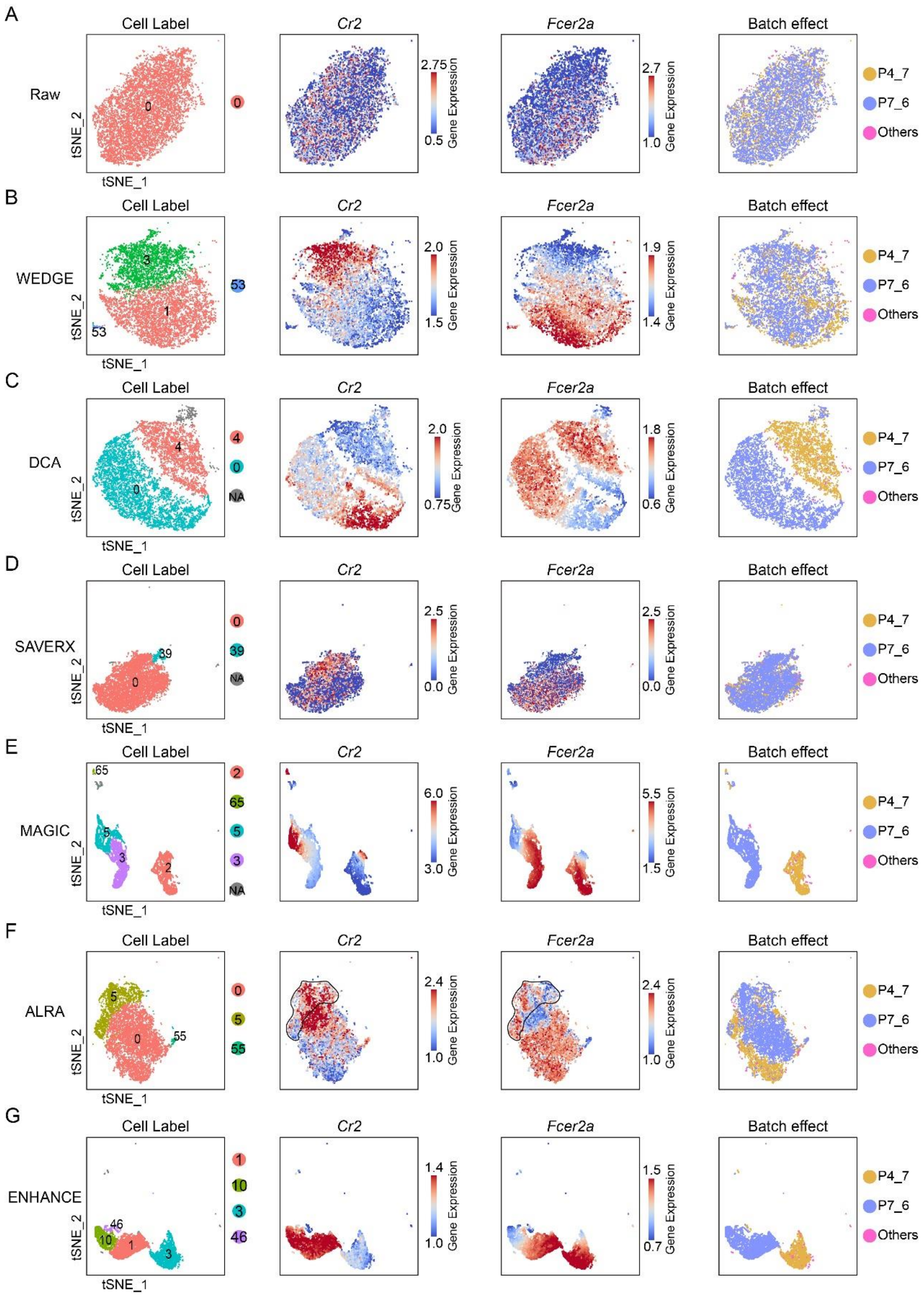


**Supplemental Figure S9.** The comparison between WEDGE and other state-of-the-art imputation methods in terms of discovering B cell subpopulations from Tabula Muris dataset. (**A**) The clustering result and marker gene expression values of the splenic B cells generated from the raw data. From the left panel to the right panel, the colors indicate cell clusters, *Cr2* expression, *Fcer2* expression, and experimental batches. (**B**-**F**) The clustering results and marker gene expression values of the splenic B cells generated by different methods, including WEDGE (B), DCA (C), SAVER-X (D), MAGIC (E), ALRA (F), and ENHANCE (G). The color scheme is the same as in (A).

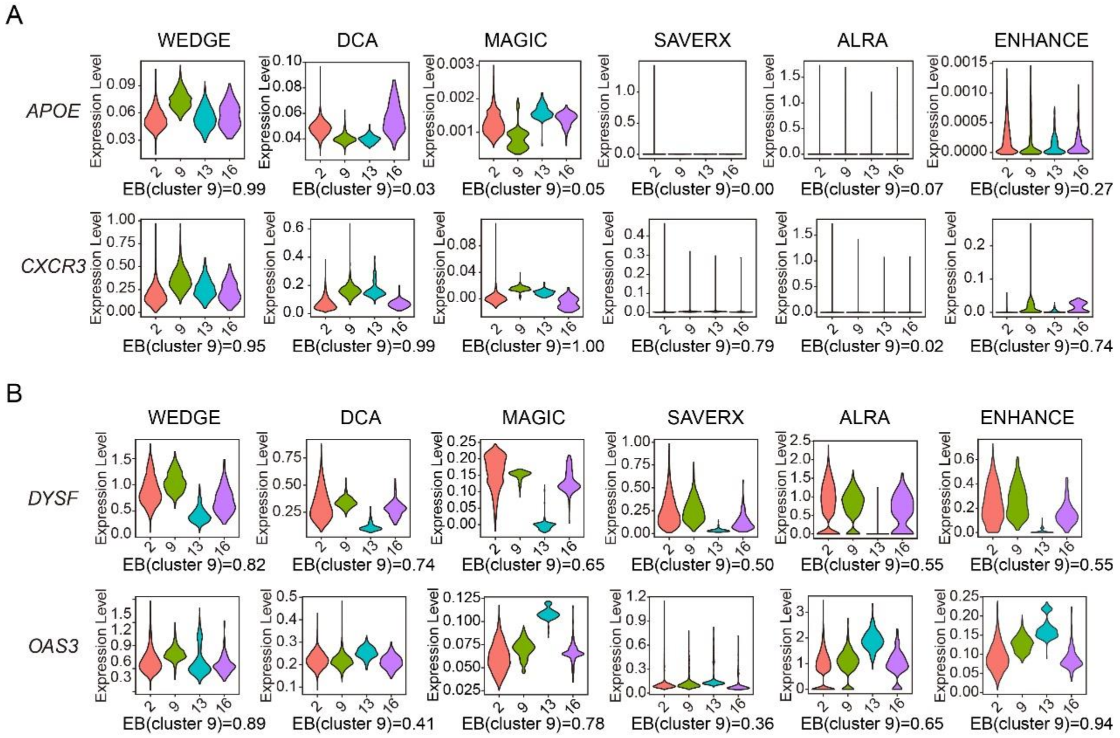

**Supplemental Figure S10.** The performance of different imputation methods on enhancing the expression of severe-stage-specific DE genes. (**A**) Inflammation-related marker genes reported by Guo *et al.* [30]. (**B**) Highly expressed genes in severe-stage COVID-19 patients reported by Wilk *et al.* [38]. The results of other methods are not shown, as SCRABBLE cannot finish the imputation in 100 hours on the computer with 72 CPU-cores (2.2GHz) and 1TB memory, and VIPER and netNMF-sc reported memory errors.

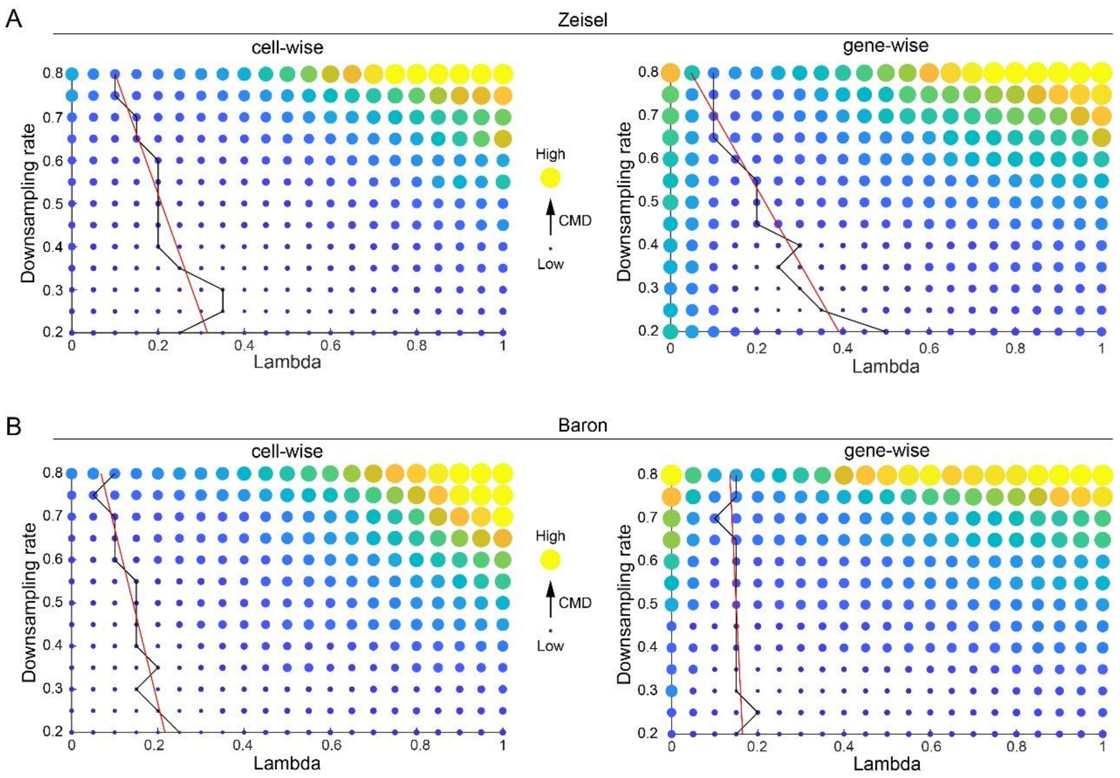


**Supplemental Figure S11.** The recovery errors (i.e. CMD) of WEDGE when using different bias parameter λ for single-cell data with different dropout rates. (**A**) The CMD values of the recovered cell-cell (left panel) and gene-gene (right panel) correlation matrices, for the data down-sampled from Zeisel's dataset. The black polyline connects the lowest CMD values of rows, and the red straight line represents the linear fit of these points. (**B**) The CMD values of the correlation matrices for the data down-sampled from Baron's dataset, as in (A).
